# Identification of flowering-time genes in mast flowering plants using *de novo* transcriptomic analysis

**DOI:** 10.1101/613745

**Authors:** Samarth, Robyn Lee, Jason Song, Richard Macknight, Paula E. Jameson

## Abstract

Mast flowering is synchronised highly variable flowering by a population of perennial plants over a wide geographical area. High seeding years are seen as a threat to native and endangered species due to high predator density caused by the abundance of seed. An understanding of the molecular pathways that influence masting behaviour in plants could provide better prediction of a forthcoming masting season and enable conservation strategies to be deployed. In this study, a high-throughput large-scale RNA-sequencing was performed on two masting plant species, *Celmisia lyallii* (Asteraceae), and *Chionochloa pallens* (Poaceae) to develop a reference transcriptome for functional and molecular analysis. An average total of 33 million 150 base-paired reads, for both species, were assembled using the Trinity pipeline, resulting in 151,803 and 348,649 transcripts respectively for *Celmisia* and *Chionochloa*. The two datasets generated were blasted against the publicly available databases, TAIR, Swiss-Prot, non-redundant protein (nr), KEGG and COG for unigene annotations. On average, 56% of the unigenes were finally annotated with gene descriptions mapped to known protein sequences for both the species. Gene ontology analysis was then performed on the assembled reference transcriptomes, categorising the transcripts on the basis of putative biological processes, molecular function, and cellular localisation. A total of 543 transcripts from *Celmisia* and 470 transcripts from *Chionochloa* were also mapped to unique flowering-time proteins identified in Arabidopsis, suggesting the conservation of the flowering network in these wild alpine plants, growing in natural field conditions. These genes can further be analysed to understand the molecular regulation of the reproductive phase transition in the masting plants.

## Introduction

RNA-sequencing (RNA-seq) has enabled the expansion of molecular biology into the fields of ecology and evolution (1). Compared to microarrays, it is the preferred high-throughput method in terms of time and cost. Researchers can analyse not only differentially expressed genes but also identify splice variants, rare transcripts, and novel small RNA targets in plants (2–4). RNA-seq has been gaining in popularity, especially in non-model plant species where sequencing and assembling large genomic datasets is complex and expensive (5). A search of the Web of Science (Thomson Reuters) database with the term RNA-seq/RNAseq resulted in 2,147 articles published between 2010-2019 with 114 articles already published in 2019 in the field of plant sciences (Fig. S1). Refining the search to the field of publication from ecology and evolution returned 734 articles published in the last five years. The application of next-generation sequencing has led to the discovery of novel genes and associated molecular markers from non-model plants which have been used to improve breeding efficiency, quality and nutritional value of agronomic crops (6, 7).

RNA-seq is responsible for recent advances in integrative biology directed at understanding the molecular mechanisms underlying behavioural, adaptive and phenotypic plasticity in wild populations. It enables characterization of the molecular nature of ecologically important traits associated with inter-individual or inter-population variation, such as disease resistance, mating behaviour, and stress-responsive genes, which mitigate the impact of changing environmental conditions (5).

A defined flowering time is a critical ecological trait necessary for a plant to achieve its reproductive potential (8). Flowering at the appropriate time enables greater seed set and distribution, and the avoidance of abiotic or biotic stresses prevailing in the environment (9). At the population level, the timing can affect the fitness of the individual species along with the prevailing competition in an ecosystem (10). Synchronisation of flowering with the growth and development cycles of insects and animal pollinators will allow greater pollination success which, in turn, will affect the species distribution as well as species richness in a geographical area (11).

Researchers have used high-throughput RNA-seq to study flowering time in perennial plants grown in natural conditions. They have identified essential genes associated with the perenniality of plants and their responses to seasonal changes involving temperature and day length (12). Flowering in perennials is a continuous process of alternation between vegetative and reproductive phases, unlike annual plants which reproduce once and die (13). On the other hand, some species show supra-annual synchronous flowering within a plant population at irregular intervals. These plant species are known as masting plants with high fluctuating seed production ability over years (14). This unique phenomenon has supra-annual patterns caused by changes in seasonal temperatures over two successive years as hypothesized from long-term phenological data (15). Evolution of masting in a plant species is a compelling study to understand whether it is a selectively favoured phenomenon or an ecological phenomenon that requires specialized physiology. Masting plants have adopted such behaviour with an inescapable energy cost including the delayed chance of reproduction and a high density-dependent mortality rate (16). The ability of the plant to sense the environmental cues to attain variability in their physiological responses even at the population level adds even more complexity.

Masting provides selective benefits to the plant population involving predator satiation, greater pollination efficiency, better seed dispersal, and food for native birds and mammals (17). However, it also leads to an explosive increase in the population of predatory animals feeding on endangered and endemic birds in Aotearoa New Zealand. Masting events are responsible for significant decreases in the population of kiwi, kakapo, kea due to the attacks by huge population of rats and stoats (14, 18). Therefore, understanding the molecular causative mechanism of a masting event can empower the phenological and mathematical datasets for systematic conservation planning of natural ecosystem.

Synthesising the molecular knowledge about masting can also benefit the ecological and evolutionary studies to draw the inferences from the phenological data available from diverse geographical habitats. Enrichment of molecular knowledge can allow forecasting future changes in the masting behaviour and assess how future changes in the natural conditions will lead to molecular adaptation of flowering-time genes and associated regulatory mechanisms affecting the masting pattern. In this study, we used RNA-sequencing and gene expression analysis to identify the potential flowering-time genes which may be involved in mast flowering in *Chionochloa pallens* and *Celmisia lyallii*. *Celmisia lyallii* (Asteraceae) and *Chionochloa pallens* (Poaceae) are two alpine perennial plants exhibiting strong masting behaviour (19). Expression analyses of the flowering-time marker genes may help to unravel the threshold and accurate timing of the reception of the flowering cue in plants associated with natural environmental conditions.

## Materials and methods

### Plant material and study sites

Samples were collected from the natural site of *Chionochloa pallens* at 1070 m and *Celmisia lyallii* at 1350 m of Mt Hutt (43.4717° S, 171.5264° E) during late summer. The Department of Conservation permit number 40225-FLO allowed for the collection of the plants used in this study. Three replicates from four different tissues (roots, mature, old and young leaves) from *Celmisia* were collected for RNA-sequencing. For *Chionochloa*, three independent replicates each comprising of three separate leaf and apical meristem tissue were collected for sequencing analysis. Leaf tissue samples were also collected for both the species from their respective field site throughout the year. The samples were stored in −80 °C until further processing.

### RNA extraction and *de novo assembly*

Total RNA was extracted from the *Celmisia* (leaves and root samples) and *Chionochloa* samples (leaf and meristematic tissue) using Qiagen Plant RNA extraction kit. The extracted RNA was DNase I digested and checked for quality using a Bioanalyser 2100 (Agilent Technologies). RNA isolated from each of the samples had a RIN of greater than 7 and rRNA ratio of greater than 1, both being essential parameters for RNA quality prior to sequencing. cDNA libraries were prepared using Illumina TruSeq kit 2.0 followed by paired-end sequencing on the Illumina HiSeq 2500 platform (20). Sequencing for *Celmisia* was done at Macrogen Inc. (South Korea), while sequencing for *Chionochloa* was carried out at NZGL (Dunedin, New Zealand). The isolated RNA from the leaf tissues and roots of *Celmisia* were pooled together to generate one cDNA library. For *Chionochloa*, separate libraries were prepared for the three biological replicates of leaf and apical meristem tissue.

The 150–base pair paired-end reads that were generated were checked for quality. Bases with a quality score of less than 30 and reads containing fewer than 20 bases were removed using the fastq quality trimmer program. Adapters were removed using the Trimmomatic program with default parameters (21). The quality of the raw data generated from both species was again analysed using the Fastqc program (22). The cleaned reads were then assembled together into a single reference transcriptome assembly using the Trinity pipeline with default parameters (23). The combined *de novo* assembly is referred to as the ‘*Celmisia* draft transcriptome’ and ‘*Chionochloa* draft transcriptome’. These transcriptomic datasets were used in all the analyses reported in this paper. All the sequences in these datasets have been deposited at NCBI as sequence read archive files [the accession numbers will be provided upon acceptance]. The Trinitystats.pl script was used to assess the completeness of the assembled transcriptome based on the assembly statistics. Additionally, the eukaryotic Benchmarking Universal Single-copy Orthologs (BUSCOs) dataset was compared with our assembled transcriptome datasets to identify single gene copy orthologous sequences (24).

### Functional annotation of the assembled transcriptome

Open Reading Frames (ORFs) with a length of at least 100 amino acids were predicted using the Transdecoder script from the assembled transcriptomes. Standalone BLAST was used to perform sequence similarity searches to annotate the obtained unigenes. The transcriptomic assembly was searched against publicly available genomic databases of *Arabidopsis thaliana* (arabidopsis), *Helianthus annuus* (sunflower), and *Chrysanthemum* (chrysanthemum) for *Celmisia*, and arabidopsis, *Zea mays* (maize), *Orysa sativa* (rice) for *Chionochloa* using blastx with an E-value cut off of 10^−3^. The predicted peptides annotated from the blastx searches were again filtered using a cut-off value of 50% identity match and 60% query coverage for stringent gene annotation. The unique gene identifiers obtained from the similarity search were then used for Gene Ontology (GO) analysis. GO analysis was performed using PANTHER based on the search output from the BLAST results to classify unigenes into molecular function, biological processes, and cellular component categories (25). The unigenes obtained were also blasted against the NR, Swiss-Prot, PlantTF databases and COG (E-value ≤ 1.0E-5) to further annotate the unigenes. Unigenes aligned to the COG database were categorised into their possible functions. Unigenes were also mapped to the KEGG database with an E-value threshold of 10^−5^ to explore pathway mapping and function in both species (26).

### Differential expression analysis

The transcript abundance estimation was performed between leaf and apical meristem tissue from *Chionochloa* using Trinity-inbuilt script. A transcript abundance matrix was generated using bowtie 2.0 by aligning the trimmed reads from each sample to the ‘*Chionochloa* draft transcriptome’. The DESeq 2 package in R was used to calculate the differential transcript expression between different samples based on the generated transcript abundance matrix (27). The differentially expressed unigenes were mapped against the Flowering-Interactive Database (FLOR-ID) to identify floral genes which may be involved in the activation of the flowering process in *Chionochloa*.

### Identification of flowering-time genes

The assembled transcriptomes were imported and maintained in the form of a collective database on the CLC genomics workbench and are referred to as the ‘*Celmisia* database’ and ‘*Chionochloa* database’ respectively. Flowering-time gene sequences from the FLOR-ID database of arabidopsis were used as the reference sequences for identification of the corresponding homologs in *Celmisia* and *Chionochloa*. The assembled transcriptomes were BLAST searched against the FLOR-ID (28) database with E-value 1e-5 and 70% protein identity. Additional, similar homologous protein sequences of flowering-time genes from sunflower and rice were also BLAST searched against the assembled transcriptomes for accurate characterisation of flowering-time genes in the *Celmisia* and *Chionochloa* databases using tblastn. The sequence with the highest score and lowest e-value was selected as a putative target sequence along with the results from the search against the arabidopsis flowering database.

### Reverse transcriptase quantitative PCR (RT-qPCR)

Samples collected throughout the year from the field sites were used to analyse seasonal expression of selected flowering-time genes. Validation of the differential expression analysis was also done using the RT-qPCR. Total RNA was isolated using a Plant RNA extraction kit from Qiagen as mentioned above. cDNA was synthesized using 1 µg of RNA and home-made reverse transcription mix. Real-time quantitative PCR was carried out using 15 µL SYBR reaction mixture (Kapa Biosystems) in a RotorGene Q cycler (Qiagen). Relative gene expression levels were calculated using the 2^−ΔΔCt^ method (29, 30). No-template controls and negative reverse transcriptase reactions were also set up alongside the qPCR batch to confirm the absence of genomic DNA and other contaminants in the sample. Protein pyrophosphatase 2A and GAPDH for *Celmisia*, Expressed protein and Tumour homolog protein for *Chionochloa* were used as the reference genes. The list of primer sequences used in the present investigation is given in supplementary S2 table.

## Results and discussion

### RNA extraction and *de novo* transcriptome assembly

The cDNA libraries, developed from leaves and root samples of *Celmisia* and leaves and apical meristem samples of *Chionochloa* and sequenced using Illumina HiSeq, generated 37,147,626 and 29,282,017 reads for *Celmisia* and *Chionochloa*, respectively. Sequencing reads were then analysed for an initial quality check to remove adapters and ambiguous bases using Trimmomatic v3.0. For *Celmisia* and *Chionochloa*, 37,030,237 and 29,281,644 cleaned reads, respectively, were used for generating the transcriptome assembly using the Trinity pipeline. The *Celmisia* and *Chionochloa* draft transcriptomes consisted of about 151,803 and 348, 649 transcripts, respectively. The average median contig length for the *Celmisia* assembly was 780 with N50 = 1524, while the median contig length for the *Chionochloa* assembly was 506 with N50 = 795. The total assembled transcriptomic size for *Celmisia* and *Chionochloa* were 164 Mbp and 245 Mbp, respectively (Table 1). The assembled transcriptomes were used in all further analyses. From these assembled transcriptomes, 136,182 and 346,249 transcripts were predicted to encode peptides which were at least 100 amino acids long for *Celmisia* and *Chionochloa*, respectively.

**Table 1:**
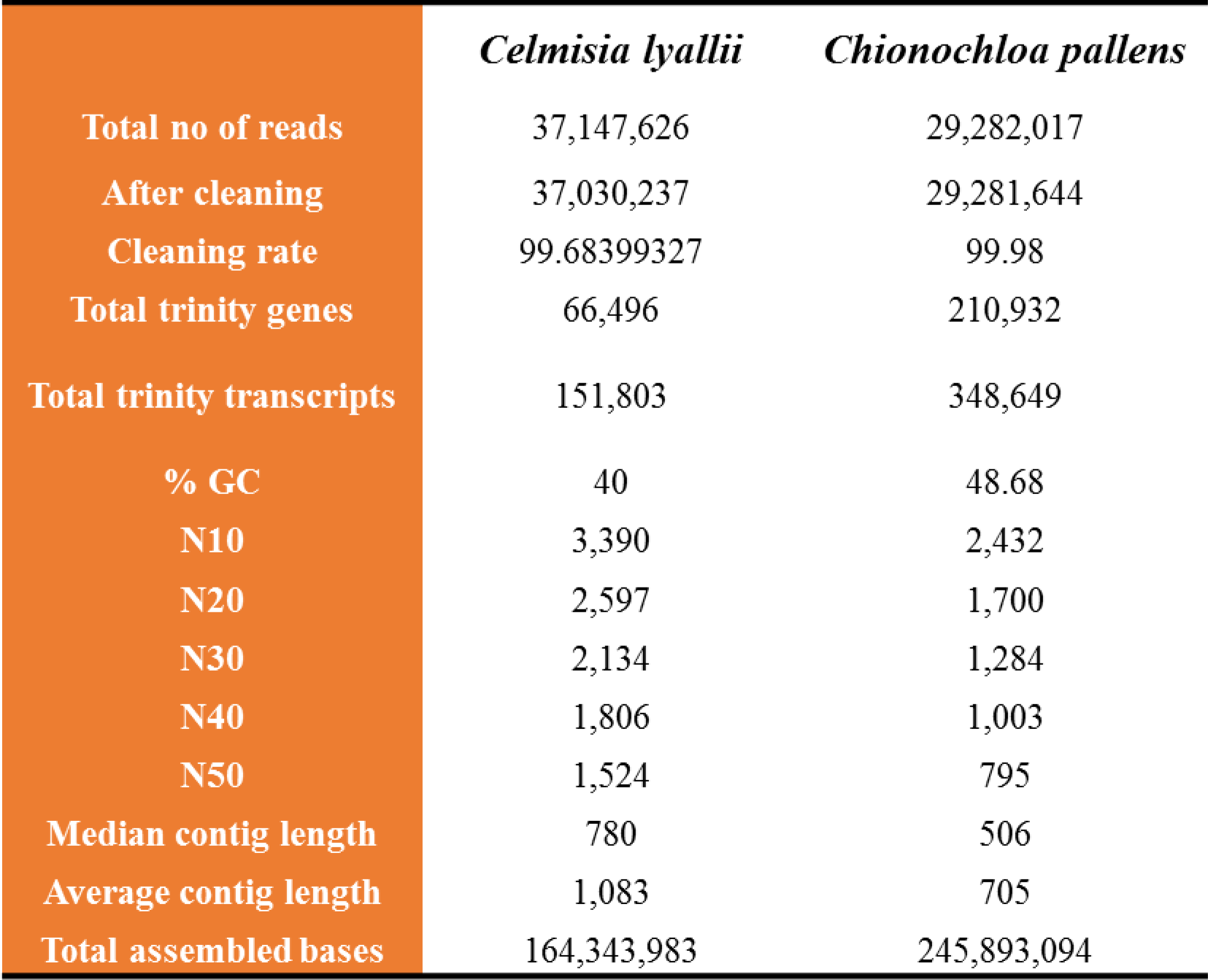
Assembly statistics for the *de novo* transcriptome

|  | *Celmisia lyallii* | *Chionochloa pallens* |
| --- | --- | --- |
| Total no of reads | 37,147,626 | 29,282,017 |
| After cleaning | 37,030,237 | 29,281,644 |
| Cleaning rate | 99.68399327 | 99.98 |
| Total trinity genes | 66,496 | 210,932 |
| Total trinity transcripts | 151,803 | 348,649 |
| % GC | 40 | 48.68 |
| N10 | 3,390 | 2,432 |
| N20 | 2,597 | 1,700 |
| N30 | 2,134 | 1,284 |
| N40 | 1,806 | 1,003 |
| N50 | 1,524 | 795 |
| Median contig length | 780 | 506 |
| Average contig length | 1,083 | 705 |
| Total assembled bases | 164,343,983 | 245,893,094 |

Many genomic features, including gene length, gene density, meiotic recombination and gene expression, have often been shown to be associated with the GC content of an organism. The GC content is highly variable among distinct species and is usually more pronounced in heterogeneous populations (31). Eukaryotes have a GC content varying between 20 and 60%. (32). The draft transcriptome of *Celmisia lyallii* has a 40% GC content which is consistent with the other dicot plant species (33). In contrast, *Chionochloa pallens* has 48% GC content suggesting it may be a wild ancestral member of the Poaceae family (34). A high GC content is often associated with high tolerance/adaptive capability. This could explain why *Chionochloa pallens* is widespread across New Zealand and adapted to varied environmental conditions.

### BUSCO analysis

Benchmarking Universal Single Copy Orthologs (BUSCO) identifies COG genes from the assembled transcriptome to identify the orthologous core-genes present in the BUSCO pipeline (24). BUSCO acts as an assessment of the transcriptome assembly by measuring the completeness of the transcriptome based on evolutionary present universal single-copy orthologs. The search revealed 87% of complete BUSCOS and 6% of fragmented BUSCOs indicating a near complete transcriptome for *Celmisia*. Similarly, 64% of complete BUSCOs and 9% of fragmented BUSCOs were found in case of *Chionochloa*, suggesting a good transcriptome assembly.

### Functional annotation of the assembled unigenes

All the unigenes from both species were functionally annotated using BLAST platform against the proteomes of multiple species. By using a combined annotation from multiple species, the power of the annotation is increased and, consequently, provides a more accurate functional characterisation of the predicted peptides.

In total, 95,993 (70%) of the *Celmisia* transcripts were annotated with 36,595 unique protein IDs. Most of the protein sequences were predicted by the NR database. The greatest number of predicted homologous sequences belong to *Helianthus annuus* followed by *Chrysanthemum sp*, both belonging to the Asteraceae family. 90% of the translated amino acid sequences of the predicted transcripts were similar to the NR database (ranging between 60% to 100% amino acid similarities). The predicted protein sequences were also BLAST searched against the proteome data from sunflower and Chrysanthemum, separately. A total of 78,249 transcripts were annotated from the combined proteomic datasets of arabidopsis, sunflower and chrysanthemum. About 58% and 50% of the total predicted protein sequences in *Celmisia* were successfully annotated from the two reference proteome datasets of sunflower and chrysanthemum, respectively, suggesting strong conservation of genetic networks between the members of the same family. BLAST searches against the uniprot database showed 53.89% similarity to the assembled *Celmisia* transcriptome, with an E-value less than 1E-5.

A BLAST search against the NR protein database yielded annotation of 160,482 (46%) transcripts present in the assembled draft transcriptome of *Chionochloa*. Transcripts were also BLAST searched (using blastp) against the proteomic datasets of arabidopsis, rice and maize to obtain gene annotations. A total of 135,372 transcripts were annotated using the combined proteomic datasets. Most of the *Chionochloa* transcripts (94,616) showed significant hit similar to the maize proteins with a shared 87% of average identity. Most of these transcripts have an E-value of less than 10^−6^ and more than 90% query coverage. About 42,235 and 89,299 unigenes were found to be significantly similar to *Arabidopsis* and rice proteins, respectively. Similarly, 93,762 transcripts (E-value < 1E-5) showed on average 58% similarity against the uniprot database.

Each unigene in the assembled transcriptome was further categorised into three GO categories of ‘molecular function’, ‘biological process’, and ‘cellular component’ using PANTHER based on the sunflower and rice protein databases used as references for *Celmisia* and *Chionochloa*, respectively. The assembled transcripts of *Celmisia* and *Chionochloa* were grouped into 33 different functional groups. A similar categorical pattern for functional grouping was observed for the annotated unigenes of both *Celmisia* and *Chionochloa*. Eight categories were found within the ‘molecular function’ subgroup, 18 within ‘biological processes’ and seven within ‘cellular component’ functional groupings for both *Celmisia* and *Chionochloa*. The top GO terms for ‘cellular component’ belonged to the cell component followed by the organelle functional group. For ‘biological processes’, the top GO terms were binding and catalytic activity. For ‘molecular function’, the top GO terms belonged to catalytic activity followed by binding and transporter activity (Fig 1a and 1b). These functional groups are a reflection of current physiological responses and their regulation in the plants as they were growing under natural conditions.

**Fig 1a and 1b.**
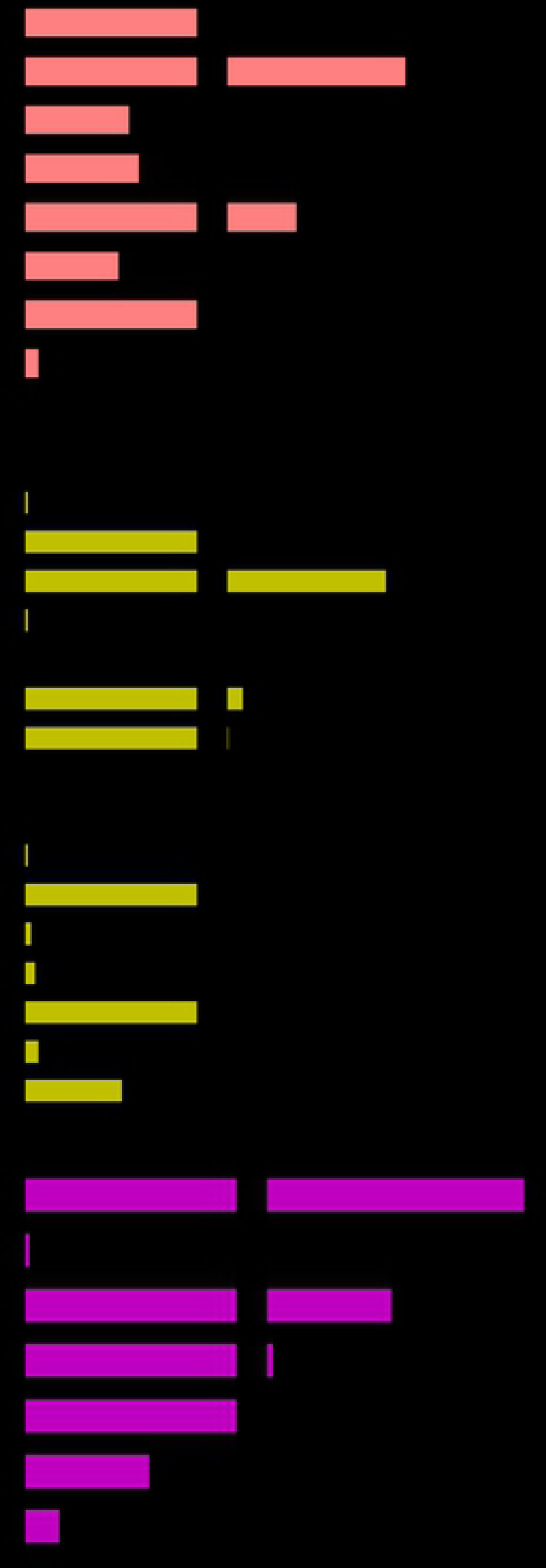
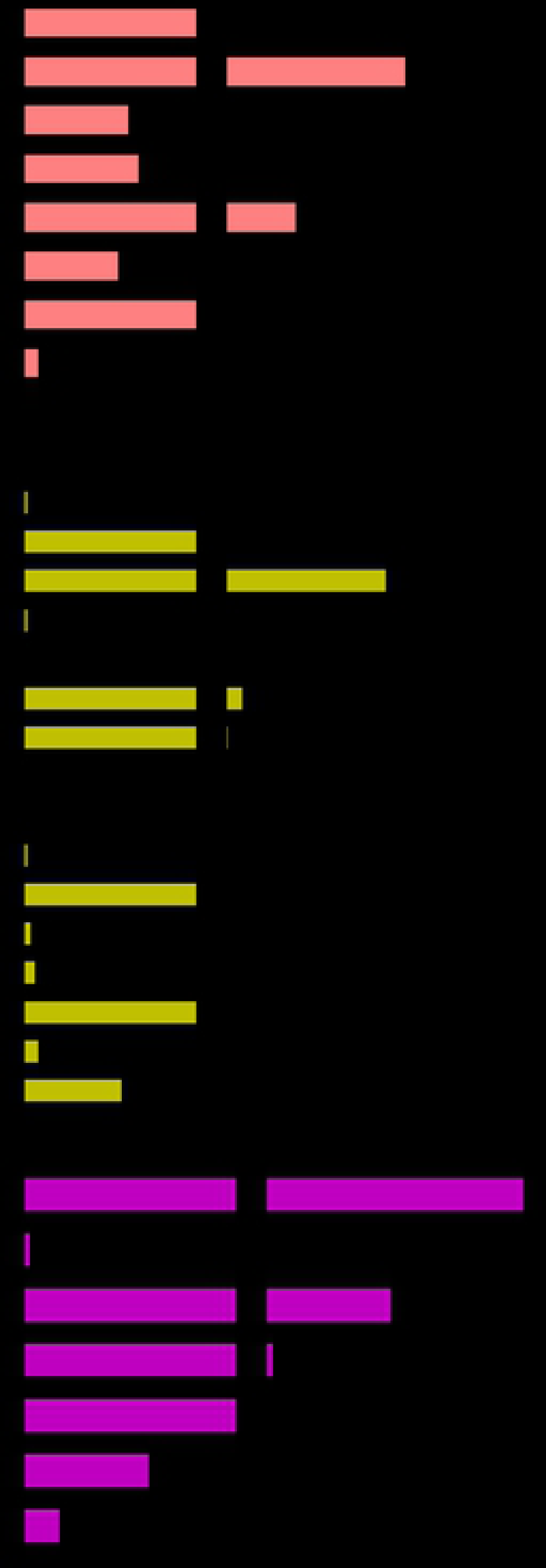
GO analysis of the assembled transcriptomes of (a) *Celmisia* (b) *Chionochloa*. Transcripts were categorised into 33 different GO categories sub-divided into molecular function, biological process, and cellular component.

### Transcription factor prediction

Transcription factors (TFs) play an important role in the regulation of growth and development. Several TF families, such as MADS-box or bHLH proteins, have been identified as key regulators of floral initiation and development. About 240 and 1696 unique transcription factors were identified from the assembled transcriptomes of *Celmisia* and *Chionochloa*, respectively (Fig 2). In *Celmisia*, the most abundant transcription factors were related to plant hormones and development, while in *Chionochloa*, VOZ-domain containing transcripts were present in abundance. VOZ transcription factors are involved in phytochrome B signalling. VOZ1 has been shown to downregulate *FLOWERING LOCUS C*, a floral repressor and to promote flowering in arabidopsis (35). The other abundant transcription factors in *Chionochloa* were also involved in growth and development processes.

**Fig 2.**
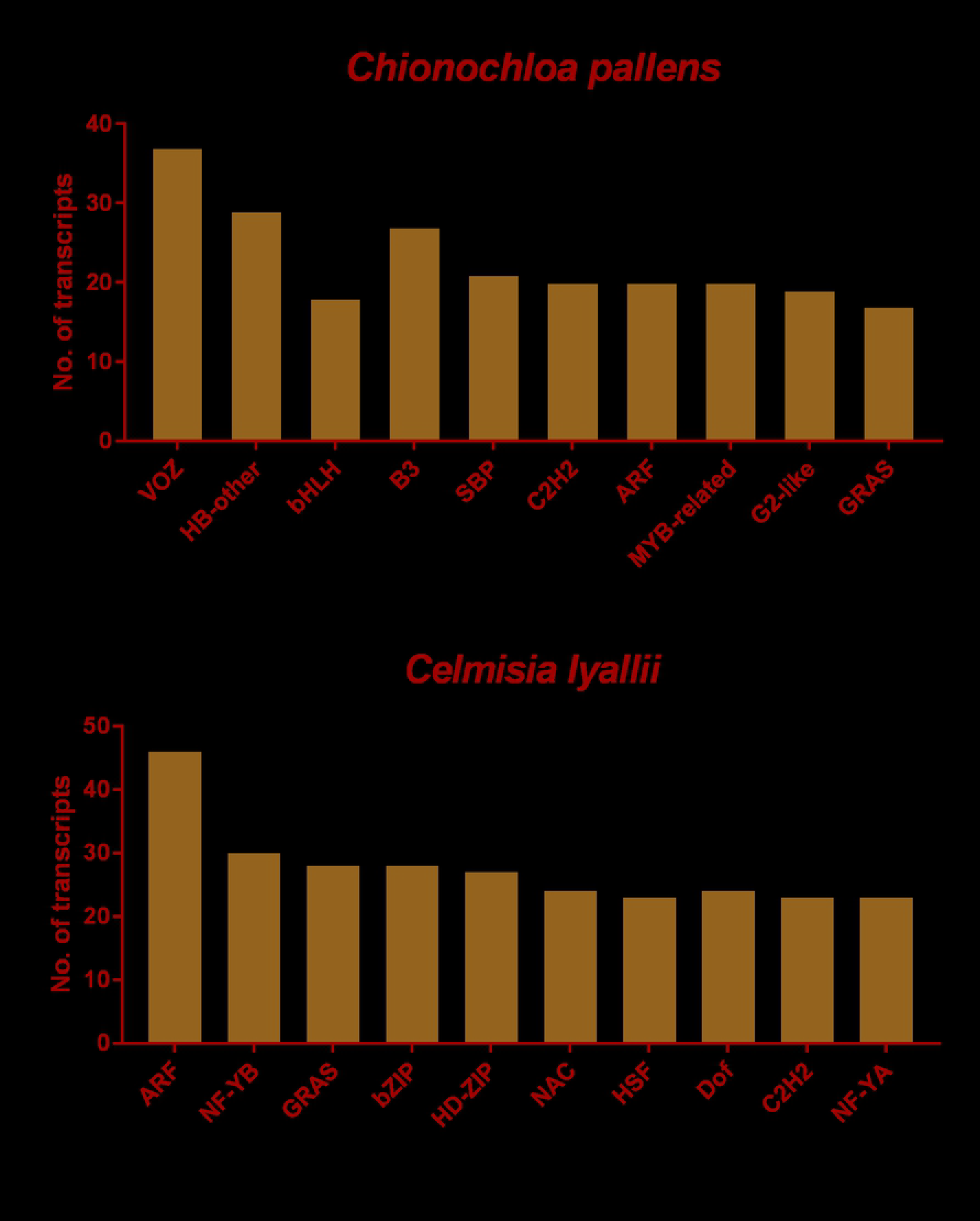
Identification of transcription factors. A subset list of the number of transcription factors identified from the assembled transcriptome of *Celmisia* and *Chionochloa*.

### COG analysis and KEGG mapping

The assembled *Celmisia* unigenes, annotated using reference proteins from *Helianthus annuus*, were further categorised into different protein classes (Fig 3). The majority of the transcripts were nucleic acid binding and hydrolase proteins, which is worth highlighting as most of the transcription factors involved in flowering are nucleic acid binding proteins. The assembled unigenes were also searched against the COG database for functional characterisation. COG is a phylogenetic database comprised of protein sequences from 66 different genomes. These COG are representative conserved domains present in either individual proteins or paralogs from three distinct lineages. Most of the proteins belonged to the general functions category followed by unknown functions and signal transduction pathways (Fig 4). A very small fraction of the unigenes were found similar to the COG database (10.2 %). Unigenes were then mapped to KEGG pathways by using the translated peptide sequences using *Arabidopsis* as a reference for pathway analysis. 26.8% of the *Celmisia* and 28.7% of the *Chionochloa* unigenes mapped to EC numbers in 119 KEGG pathways. The largest number of unigenes were mapped to metabolic pathways (11% in *Celmisia* and 12.4% in *Chionochloa*) followed by secondary metabolite synthesis (6.2% in *Celmisia* and 6.8% in *Chionochloa*).

**Fig 3.**
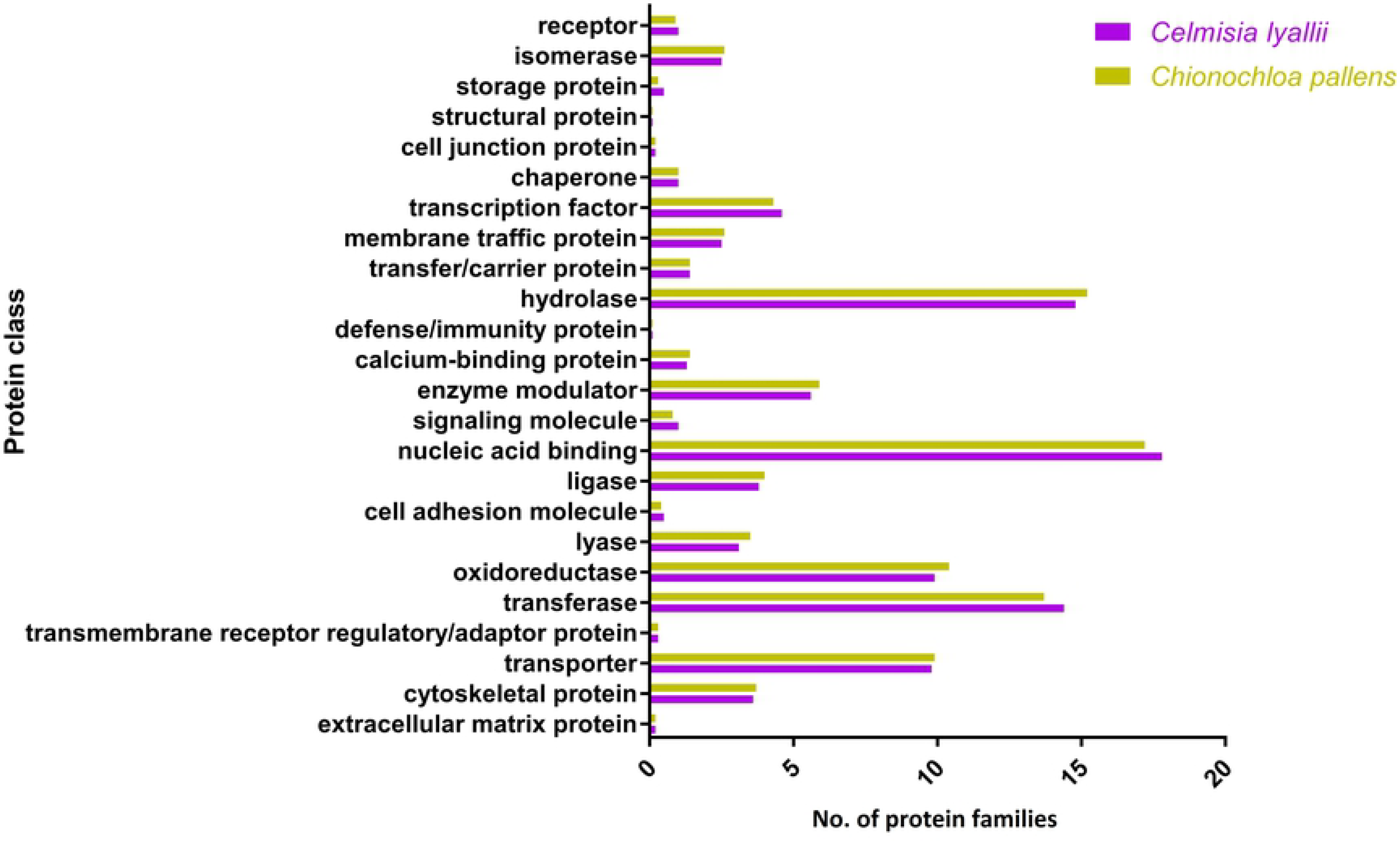
Identification of different protein classes. Categorisation of the peptide sequences predicted from the assembled transcriptome of *Celmisia* and *Chionochloa*.

**Fig 4.**
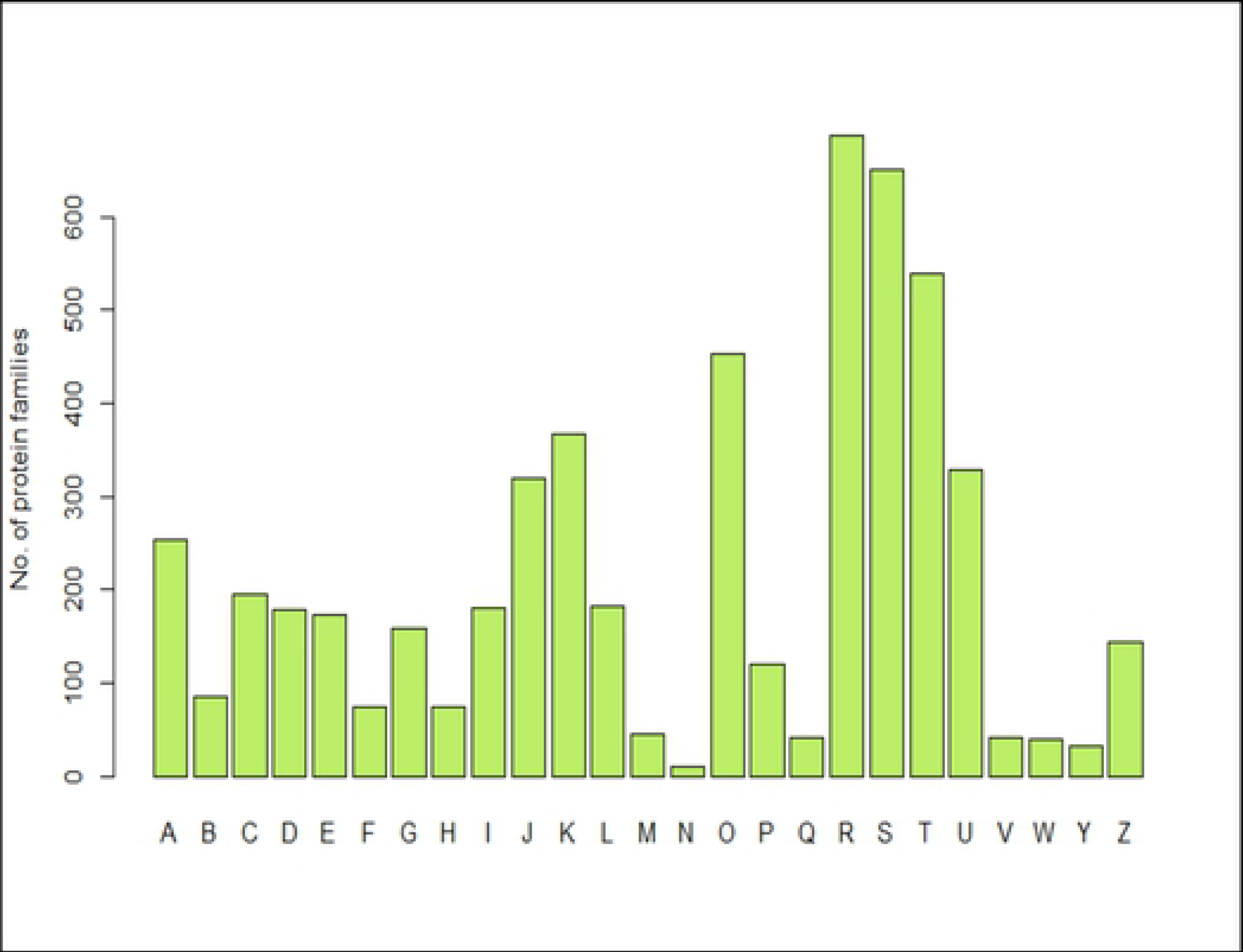
Functional classification of the *Celmisia* transcriptome. The identified unigenes were further grouped into 25 categories base on the homology search against the COG database.

### Identification of flowering-time genes in *Celmisia lyallii* and *Chionochloa pallens*

There are 306 flowering-time genes identified in *Arabidopsis* responsible for the transition to flowering. Expression analysis of these flowering-time genes may help unravel the mechanism of floral transition in masting plants using differential temperature as a cue (Kelly et al., 2013). Previously characterised flowering-time genes from arabidopsis and rice were used as references to identify the corresponding homologous sequences in *Celmisia* and *Chionochloa*, respectively. The assembled transcriptomes were imported and maintained in the form of a collective database at the CLC Genomics Workbench 8. To validate and annotate the assembled unigenes, gene sequences from arabidopsis and sunflower were used as the reference sequences for *Celmisia*. Although the core genes accountable for regulation of flowering are similar in both monocots and dicots, the flowering regulatory network consisting of temperature and photoperiodic sensing genes are different in monocots compared to dicots. Therefore, protein sequences from rice and maize were used as references for flowering gene(s) identification in *Chionochloa*, as they both belong to the Poaceae family. Using blastp from the NCBI BLAST suit, floral gene sequences from the draft transcriptomes of *Celmisia* and *Chionochloa* were identified under stringent selection criteria of E-value of less than 1E-3, 70% protein identity and 50% query coverage. Such criteria should eliminate the risk of improper identification of the homologous sequences due to phylogenetic distances and ploidy level of the genome. In total, 543 and 470 transcripts, matching with 97 and 110 unique floral protein IDs were identified in *Celmisia* and *Chionochloa*, respectively (Fig S3). These gene(s) may play a key role in coordinating the vegetative to reproductive phase transition during masting.

### Tissue-specific expression of flowering-time genes in *Chionochloa pallens*

Pairwise comparisons were used to analyse the differentially expressed genes in the sampled leaves and apical meristematic tissues of *Chionochloa*. Using a significance threshold of 0.005 False Discovery Rate and 2-fold change in expression, we determined that there were 15,324 differentially expressed genes between the two tissues (Fig 5). Transcripts showed distinct expression patterns as they emerged into two separate groups from leaves and apical meristems respectively (Fig 6). There were 124 differentially expressed transcripts belonging to 38 unique floral genes in the leaves compared to the meristematic tissue.

**Fig 5.**
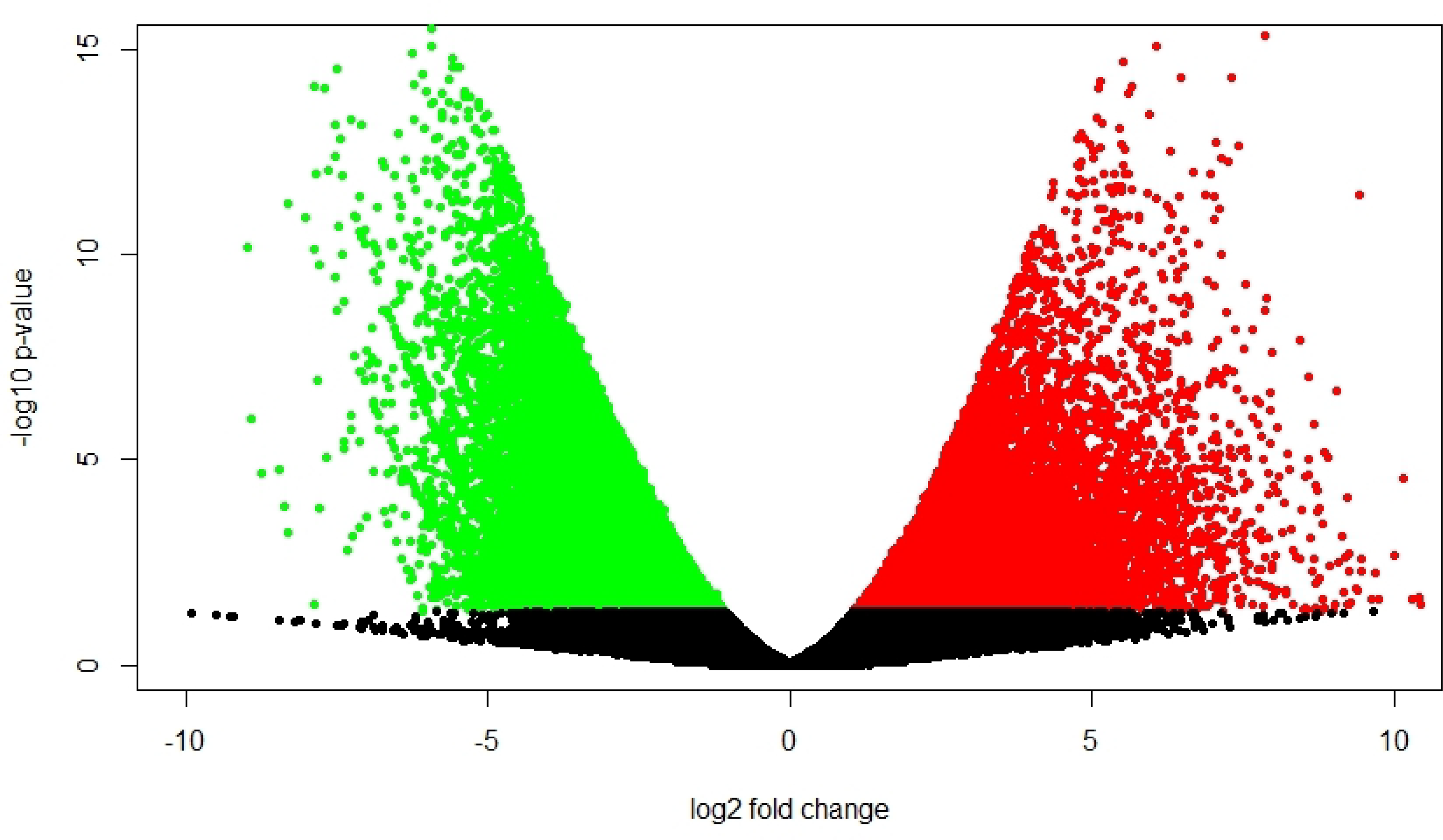
Volcano plot showing differentially expressed genes. 15,234 significantly differentially expressed genes were identified between the leaves and apical meristems. The red dots represent significantly up-regulated genes and green dots represent significantly downregulated genes with a P-value < 0.05.

**Fig 6.**
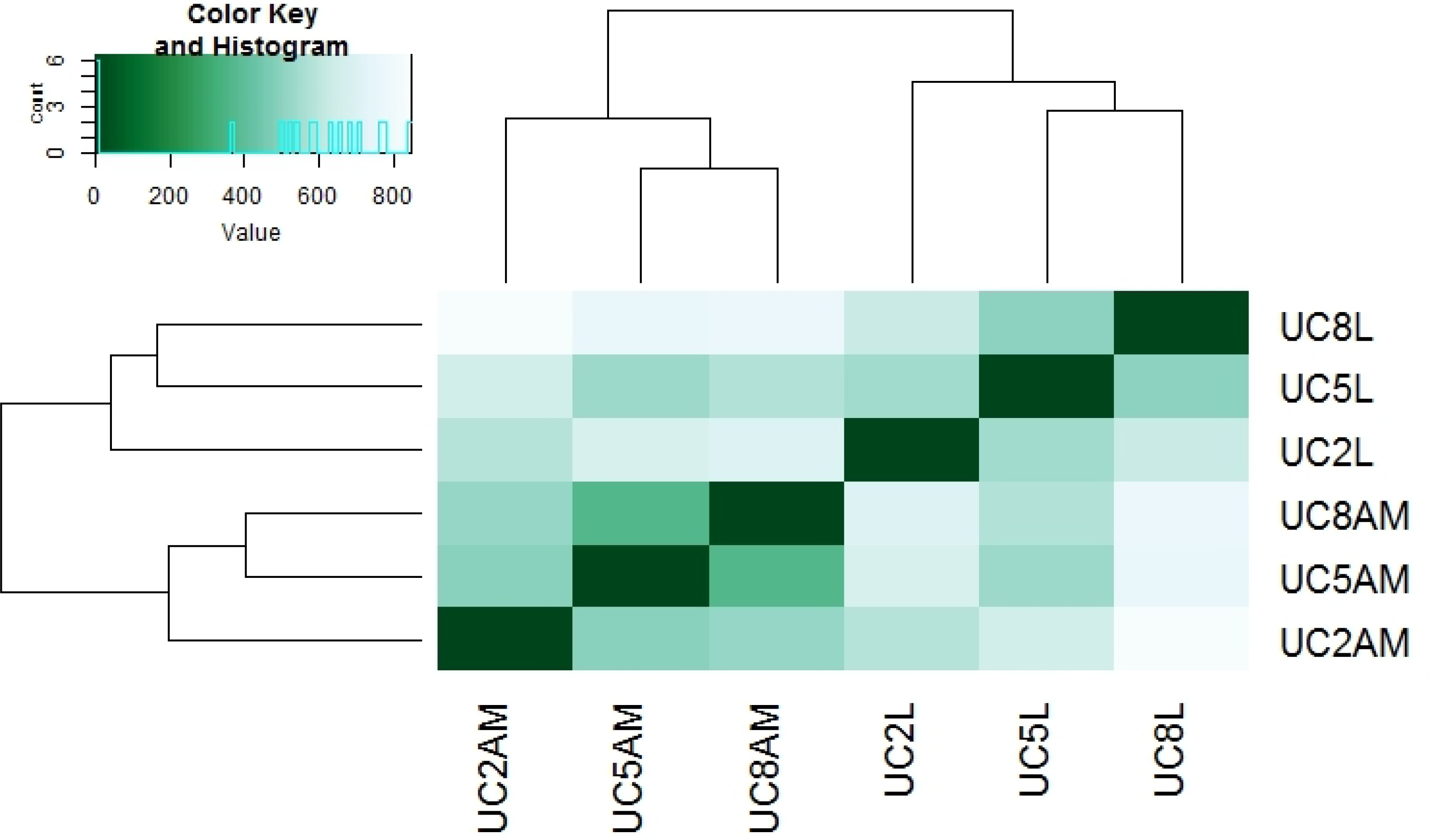
Comparisons of transcriptional profiles across samples. Heat-map showing hierarchical clustering resulting from a pairwise comparison of transcript expression levels.

Flowering is a complex molecular process regulated by several external and endogenous cues. Several activators and repressors of flowering have been characterised in arabidopsis and other plant species controlling flowering time. In the differential gene expression analysis, most of the floral activators were highly downregulated. These downregulated genes are involved in regulation of temperature, hormonal, and age-mediated flowering time control. Expression of many epigenetic modifiers responsive to temperature changes were also downregulated. Photoperiodic genes such as *CONSTANS, CRYPTOCHROME1* and *PIF4* and sugar signalling genes were upregulated in the leaves (Fig 7), changes in the expression of epigenetic modification genes and photoperiodic genes can be correlated with the presence of long days when samples were collected. This initial analysis suggests that the core components of the flowering network are conserved in *Chionochloa* and may be involved in activation of flowering during a masting event. The study also provides evidence of a role for floral epigenetic modifiers which may play a crucial role in temperature-mediated synchronised flowering.

**Fig 7.**
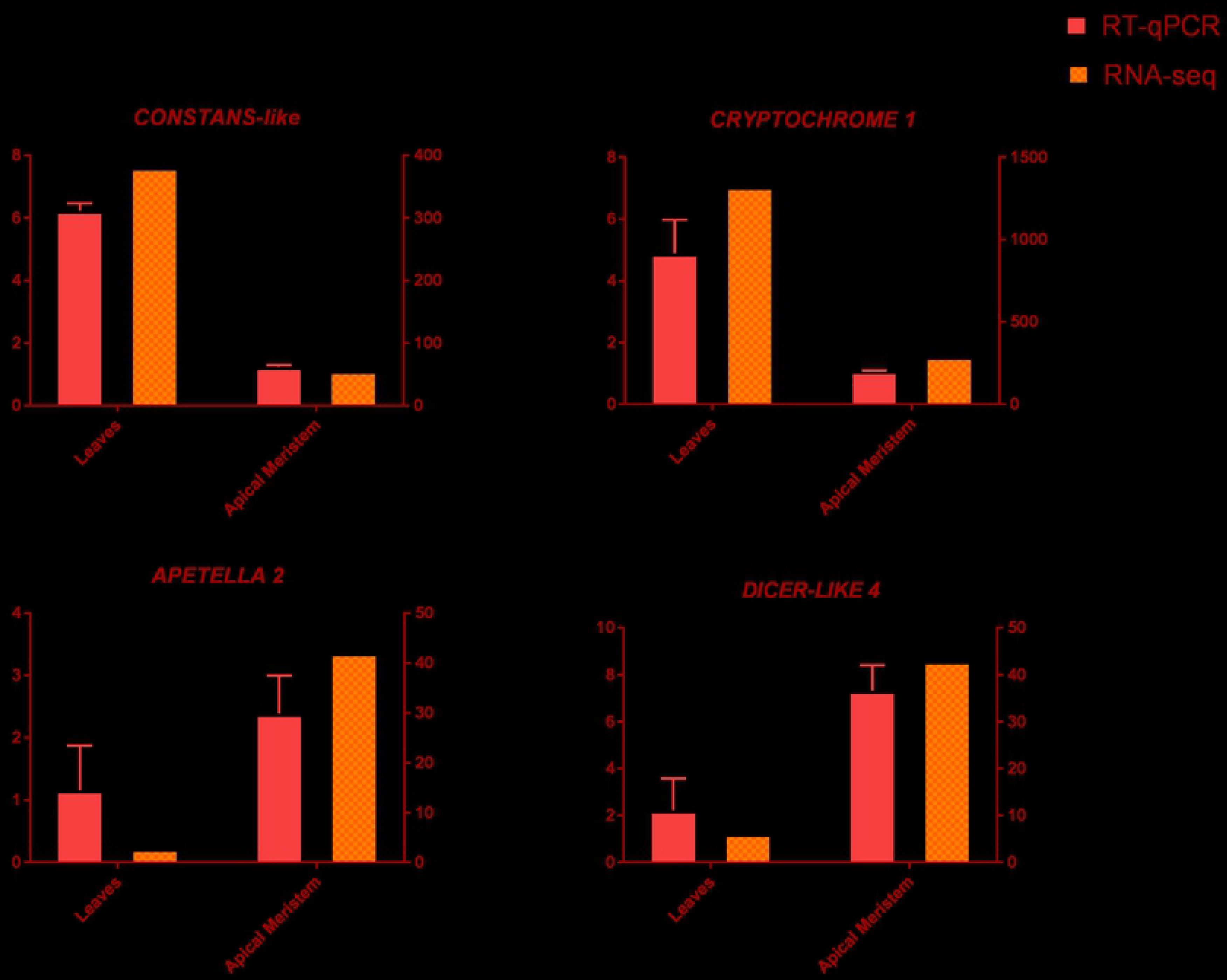
Relative expression of the flowering-pathway genes *CONSTANS-LIKE*, *CRYTOCHROME*, *APETELLA 2*, and *DICER-LIKE 4* across leaves and apical meristems. Expression profile is based on RNA-seq data, normalised by TMM method and RT-qPCR data showing relative fold change, normalised to the selected reference genes. The RT-qPCR experiment was performed using three biological replicates for each tissue with three technical replicates ±S.D.

### Seasonal expression of the flowering-time genes identified in *Chionochloa* and *Celmisia*

Expression of key flowering-time genes was monitored to determine their potential role in the growth and development of the two masting plants (Table 2). Due to destructive sampling (collection of leaves and apical meristems), it was impossible to predict whether the leaves subtended meristems that would have flowered and draw conclusions on the molecular network of flowering. Hence, leaves from both species were sampled throughout the year, representing four different seasons (summer, autumn, spring, late summer for *Celmisia* and summer, autumn, spring, winter for *Chionochloa*) (Fig 8). Expression analysis of these selected genes was carried out using RT-qPCR with two biological replicates, each of which comprised a combined pool of three individual plants.

**Table 2:** Flowering genes selected for expression analysis

| Table 2: Flowering genes selected for expression analysis |  |  |  |  |  |  |  |  |
| --- | --- | --- | --- | --- | --- | --- | --- | --- |
| *Celmisia* |  |  |  |  |  |  |  |  |
| Gene | Arabidopsis homolog | Arabidopsis locus ID | Sunflower homolog | Sunflower locus ID | % Identity | E-value | %Gaps | Contig ID |
| *Cel GI* | GIGANTEA | AT1G22770 | Ha GI | HQ428760.1 | 78.53 | 0 | 1.75 | TRINITY_DN48665_c1_g2_i1 |
| *Cel PHYB* | PHYTOCHROME B | AT2G18790.1 | Ha PHYB | GU985581.1 | 86.57 | 0 | 0 | TRINITY_DN47433_c0_g1_i1 |
| *Cel AP2* | APETALA2 | AT4G36920 | Ha AP2 | XP_021998113 | 81.65 | 1.90E-75 | 0 | TRINITY_DN45405_c1_g1_i4 |
| *Cel SVP* | SHORT VEGETATIVE PHASE | AT2G22540 | Ha SVP | XP_022001248 | 85.27 | 2.66E-96 | 0.45 | TRINITY_DN40059_c1_g1_i3 |
| *Chionochloa* |  |  |  |  |  |  |  |  |
| Gene | Arabidopsis homolog | Arabidopsis locus ID | Rice homolog | Rice locus ID | % Identity | E-value | %Gaps | Contig ID |
| *Cp GI* | GIGANTEA | AT1G22770 | Os GI | CAB56058.1 | 95.24 | 0 | 0 | TRINITY_DN35492_c2_g6_i1 |
| *Cp PHYB* | PHYTOCHROME B | AT2G18790.1 | Os PhyB | CAA40795.2 | 90.96 | 0 | 0 | TRINITY_DN28349_c2_g6_i1 |
| *Cp Ehd3* |  |  | Os Ehd 3 | BAI77463.1 | 78.35 | 3.27E-52 | 0 | TRINITY_DN32679_c1_g8_i1 |
| *Cp MADS1* | SUPPRESSOR OF OVEREXPRESSION OF CONSTANTS 1 | AT2G45660 | Maize MADS1 | NP_001105152.1 | 85.27 | 2.66E-96 | 0.45 | TRINITY_DN40059_c1_g1_i3 |

**Fig 8.**
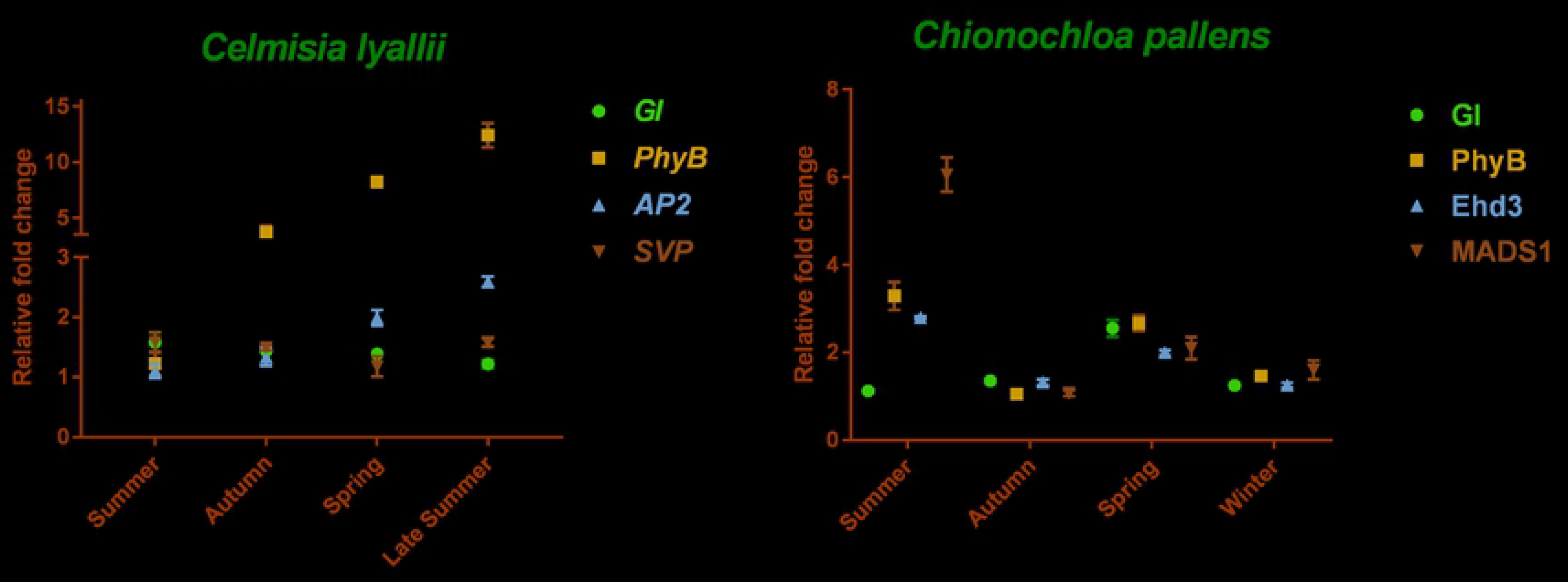
Seasonal gene expression analysis of selected flowering-pathway genes in *Celmisia* and *Chionochloa*. The relative expression was calculated using 2^−∆∆Ct^ method, represented by data from two biological replicates with three technical replicates ± S.D.

In *Celmisia*, *GIGANTEA* (*GI*), *PHYTOCHROME B* (*PhyB)*, *SHORT VEGETATIVE PHASE* (*SVP*), and *APETELA 2* (*AP2*) genes were selected for seasonal expression analysis. GI and PhyB are two crucial proteins regulating circadian rhythm and photoperiodic-mediated flowering control. *SVP* and *AP2* are regulators of age and maturity of the plant and are repressors of flowering. All the selected genes showed significant variation during the season (Fig 8). Expression of *GI* was greatest during summer, followed by a decrease in the late autumn and winter. At the same time, the expression of *PhyB* was low during the summer but increased sharply in the following seasons. The expression of *PhyB* was seven times greater in the early spring samples compared to the previous summer season. The expression pattern of *AP2* was similar to *PhyB*, with greater expression in the following spring and late summer compared to the previous summer. *SVP* had greater expression in the samples collected in summer (Fig 8). The greater expression of *SVP* during the summer season may interfere with the perception and signalling activation of the floral promoting genes. This analysis suggests that *PhyB, SVP* and *AP2* may be responsible for blocking the activation of the floral transition in *Celmisia lyallii*.

*GI*, *PhyB*, *MADS1*, and *EARLY HEADING DATE 3* (*Ehd3*) genes were selected for initial expression analysis in *Chionochloa* samples collected over the year. *MADS1* and *Ehd3* are the activators of floral promoting genes in response to temperature and day length. Expression of *GI* was found to be greater in the early spring samples compared to the other samples. There were differences in the seasonal expression pattern of *PhyB*, *Ehd3* and *MADS1*, all of these genes had a greater expression in the summer season compared to the other seasons. There was a decrease in the expression of these genes in late autumn followed by an increase in the spring. There was a high expression of *MADS1* in the summer season. However, no flowering was seen in any of the plants in the next season from which the samples were collected even though samples had a greater expression of *Ehd3* and *MADS1* in the activation period (Fig 8). The results indicate that there is another level of regulatory networks in *Chionochloa* controlling the floral transition.

## Conclusion

Ecological transcriptomes are the new means of investigating the biological processes connecting ecological concepts and molecular mechanisms. This new era has seen extensive advancement in the field of next-generation sequencing analysis. Now researchers are using these tools in the natural conditions to answer the bigger ecological questions. This study stands as a prime example of the use of ecological transcriptomes to understand masting – the intermittent synchronised flowering in perennial plants. Due to the large genomes and high ploidy levels of perennial plants, transcriptomics stands out as an exceptional alternative for gene(s) identification instead of whole-genome sequencing. In this work, high-quality RNA-seq was used to identify and study global flowering gene expression patterns between different tissues of a masting plant and across the year. These initial flowering-time seasonal expression analyses established the conservation of the flowering network in plants. The study also indicated a role for changing seasonal conditions, such as temperature or day length, in controlling the floral transition in the masting plants. Such molecular phenological studies can help in unravelling the molecular regulators of masting and explain why plants have adopted such mode of delayed reproduction. The global expression analysis between different tissues also strengthens the use of RNA-seq to study the role of summer temperatures in the occurrence of a masting event as hypothesized in Kelly et al. (2013).

## Authors’ contributions

Conceptualisation: Samarth, PEJ, RL, and RM

Data curation: Samarth and RL

Investigation: Samarth and JS

Supervision: PEJ, RL and RM

Validation: Samarth

Visualisation: Samarth

Writing-original draft preparation: Samarth and PEJ

Writing-Review and Editing: Samarth, PEJ and RM

## Acknowledgements

The authors acknowledge the University of Otago IT facility for providing the computation facilities. The authors acknowledge the Marsden Fund of the Royal Society of New Zealand for providing financial support including a Doctoral Scholarship for Samarth (Grant UOC1401). The funding body had no role in any aspect of the research. The authors also acknowledge the help from Sana Ullah Lehre for collection and preparation of plant samples for sequencing.

## Supplementary file legends

**S1 fig.** Number of RNA-seq studies published in the field of plant sciences from 2010-2019.

**S2 table.** List of primer sequences used in the present study.

**S3 table.** List of flowering genes identified in *Celmisia* and *Chionochloa* with their similarity search values.

## Data availability

The accession numbers of the data generated will be provided upon acceptance of the manuscript.

## Ethics approval and consent to participate

All plant material was collected with appropriate permissions and consultation. Department of Conservation permit number 40225-FLO allowed for the collection of the plants used in this study.

